# Immersive exposure to simulated visual hallucinations modulates high-level human cognition

**DOI:** 10.1101/2024.07.17.603874

**Authors:** Antonino Greco, Clara Rastelli, Andrea Ubaldi, Giuseppe Riva

## Abstract

Understanding altered states of consciousness induced by psychedelic drugs is crucial for advancing our knowledge of conscious perception and developing clinical applications for psychiatric conditions. Recently, technological advances in virtual reality (VR) headsets and deep neural network for generative computer vision have enabled the controlled, immersive simulation of visual hallucinations. Although there is some evidence that simulated visual hallucinations increase cognitive flexibility, comprehensive experimental data on how this artificially altered perceptual phenomenology affects high-level human cognition is lacking. We addressed this gap by measuring a wide range of behavioral tasks in human participants after the exposure to VR immersive panoramic (360°) videos and their psychedelic counterparts generated by the DeepDream algorithm. Participants exhibited reduced task-switching costs after simulated psychedelic exposure compared to naturalistic exposure when instructed to invert the stimulus-response mapping, consistent with increased cognitive flexibility. No significant differences were observed between naturalistic and simulated psychedelic exposure in linguistic association tasks at word and sentence levels. Crucially, we found that visually grounded high-level cognitive processes were modulated by exposure to simulated hallucinations, as evidenced by participants’ drawing performance. These findings reveal how altering perceptual phenomenology through simulated visual hallucinations significantly modulates high-level human cognition. Our results provide insights into the interdependence of bottom-up and top-down cognitive processes and encourage further investigation into the effects of artificial psychedelic experiences on human cognition. This research may offer valuable insights into altered states of consciousness without pharmacological intervention, potentially informing both basic neuroscience and clinical applications.

## Introduction

Psychedelics are serotoninergic drugs that profoundly alter conscious experience, by disrupting the hierarchical organization of the neural dynamics and increasing neural variability in the human brain ^1–4^. Recently, there has been a renewed interest in the application of psychedelic drugs to investigate its role in the treatment of various psychiatric and neurological conditions ^5–8^ and as a methodology to experimentally study the neural mechanisms of altered states of consciousness ^9–13^.

However, the specific contribution of these pharmacological manipulations to the altered perceptual phenomenology is difficult to estimate given that psychedelics have also secondary effects on the physiology of the human body. In addition, it is difficult to obtain approval for the use of psychedelic drugs in scientific research in many countries due to ethical and legal issues. These limitations lead to the adoption of a methodology that combined machine learning algorithms to simulate visual hallucinations with panoramic dynamic visual stimuli immersively experienced via Virtual Reality (VR) technology ^14^.

This enabled to study the phenomenological features of psychedelic altered visual perception and its effects at the behavioral and neural level, by leveraging the DeepDream algorithm ^15^ to synthesize biologically-plausible artificial hallucinations using pretrained deep convolutional neural networks (CNN) ^16,17^. With this approach, Suzuki et al. ^14^ found that DeepDream-induced perceptual phenomenology was qualitatively similar to actual psychedelic experience, probed via psilocybin, by comparing participants’ behavioral reports to a questionnaire. Then, Greco et al. ^18^ exposed human participants to similar simulated visual hallucinations in a non-immersive manner and found a similar neural dynamics between actual psychedelic experience and the simulated altered phenomenology, denoted by an increased neural variability and global functional connectivity. Following this, Rastelli et al.^19^ demonstrated that immersive simulated visual hallucinations enhanced participants’ cognitive flexibility compared to control video stimulation across various behavioral tasks.

Together, these previous studies validated the approach for simulating visual hallucinations during the psychedelic experience while providing some evidence for how cognition is affected by these alterations in the sensory domain. Rastelli et al. ^19^ particularly demonstrated how affecting low-level cognitive processes by altering visual perception influences a high-level cognitive process like cognitive flexibility. However, comprehensive experimental evidence on how this altered perceptual state via simulated visual hallucinations generally affects high-level human cognition remains lacking.

Our study addresses this research gap by measuring a wide range of behavioral tasks in human participants, covering high-level cognitive domains such as executive functions, linguistic semantic control and visual imagination. Participants were exposed to immersive (360°) panoramic videos in VR recorded in natural settings and their altered counterparts generated by DeepDream. Our results demonstrate how simulated visual hallucinations significantly modulate high-level human cognition in specific domains, providing insights about the nature of the relationship between low- and high-level cognitive processes. This research paves the way for further scientific investigations about altered states of consciousness without pharmacological intervention and suggests practical applications for modulating high-level cognition through artificially altering perceptual states.

## Results

Data were collected from 47 human participants equipped with a virtual reality (VR) headset, while they were exposed to a series of natural panoramic high-quality videos and their perceptually altered counterparts using the DeepDream algorithm (Fig. 1A). We recorded the natural videos in public places with a 360° panoramic camera, covering a wide range of ordinary situations for humans (Fig. 1A left). Perceptually altered videos were generated using two pretrained convolutional neural networks (CNN) trained on different image datasets (Fig. 1A right). During the exposure, participants were free to move their head to explore the panoramic videos. The order of presentation of conditions, which we refer as original (OR, for natural videos) and DeepDream (DD, for altered videos) to maintain continuity with previous research ^19^, was counterbalanced across participants. Immediately after the immersive exposure in each condition, participants performed four behavioral tasks, with a fixed order of execution (Fig. 1B).

**Figure 1:**
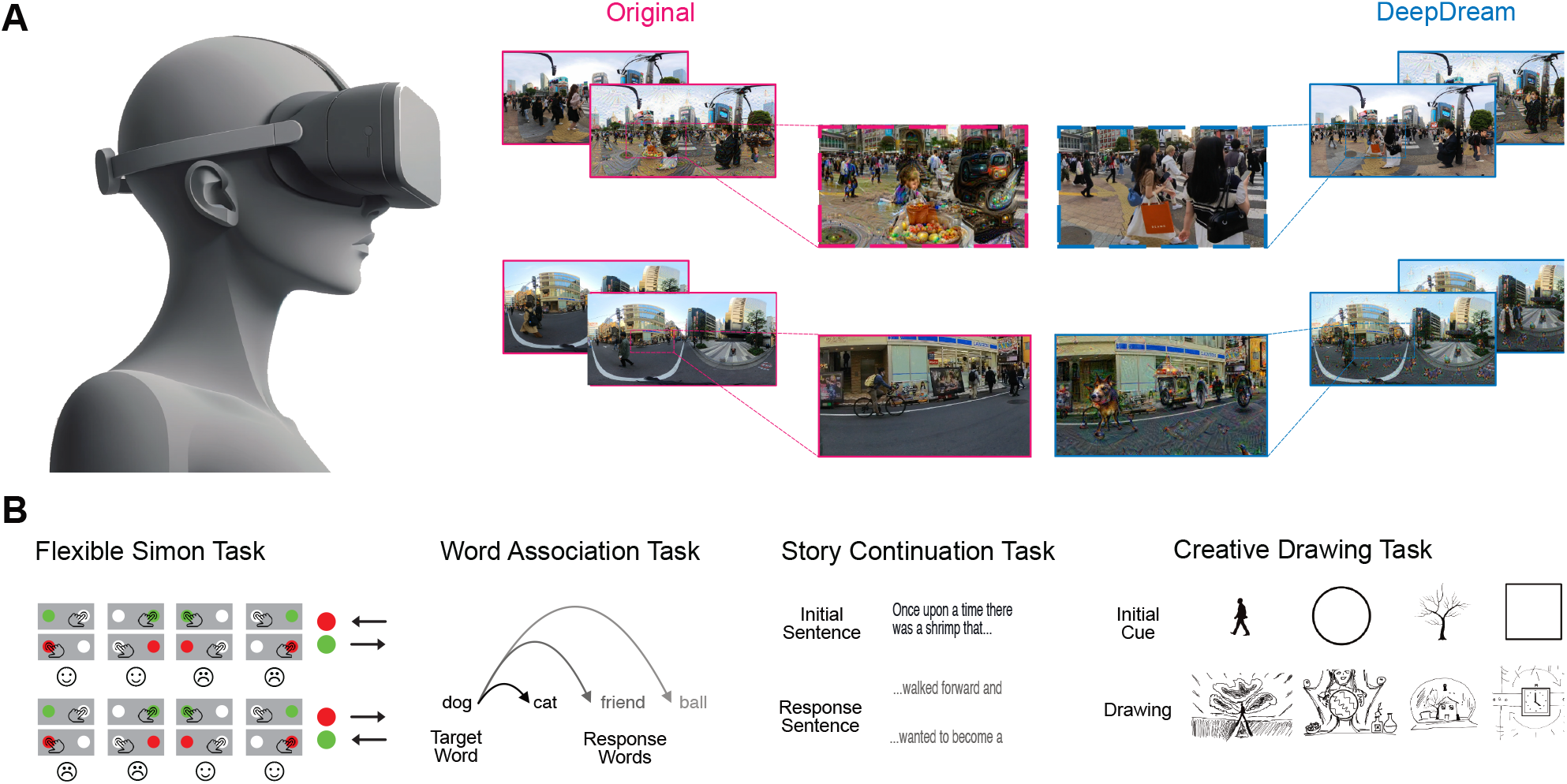
Immersive exposure to simulated visual hallucinations and behavioural tasks. **A** The experimental design comprised human participants immersed in a 360-degree panoramic video using a Virtual Reality (VR) headset as exemplified on the left. On the right, examples of frames from the video clips presented to participants in both original (pink, left side) and DeepDream (blue, right side) conditions. Top and bottom examples show the two different pretrained convolutional neural network (CNN) used to generate the psychedelic alterations to the original videos. **B** After the immersive exposure, participants performed these 4 behavioural tasks, namely the Flexible Simon Task (first to the left) investigating cognitive flexibility, the Word Association and Story Continuation Tasks (center) studying linguistic processing and the Creative Drawing Task (first to the right) investigating visually grounded high-level cognitive processes.

The first task was the Flexible Simon Task ^20^, in which participants had to map a stimulus to a motor response. In our study, they were instructed to respond with the left finger whenever they saw a red circle and with the right finger for the green circle, irrespectively of the spatial location which was either left or right (Fig. 1B left). Then, we instructed to invert the stimulus-response mapping and we computed a flexibility score by subtracting the performance of the first block from the second one (with inverted mapping) to investigate task-related switch costs ^21^. This measure allowed us to study whether the exposure to DD increased the cognitive flexibility of participants compared to OR, by comparing the accuracy and reaction times (RT) of participants across congruent and incongruent trials (Fig. 2). Crucially, we found that the flexibility score on the cumulative distribution function (CDF) of RT significantly diverged when classifying DD and OR in congruent trials (Fig. 2A, accuracy=63.8%, p=0.015). No difference was observed for CDF flexibility in incongruent trials (accuracy=51.1%, p=0.407). Thus, we tested more specifically whether the average RT flexibility score differed between conditions (Fig. 2B), founding that indeed this was the case in congruent trials (average switch costs for OR=28.6 ms and DD=4.7 ms, p=0.015, Cohen’s d=0.250) even when we consider only correct trials (average switch costs for OR=29.8 ms and DD=7.1 ms, p=0.022, d=0.236). This result showed how the exposure to DD significantly decreased the switching costs to change the stimulus-response mapping. No difference was observed for incongruent trials (p=0.893, d=0.014) even when accounting for only correct trials (p=0.535, d=0.064). We also found no difference in the accuracy flexibility score (Fig. 2B, for congruent trials p=0.662, d=0.048, for incongruent trials p=0.958, d=0.007), since the task was relatively easy to accomplish along this measure. These findings revealed how immersive exposure to simulated visual hallucinations increased the cognitive flexibility of human participants.

**Figure 2:**
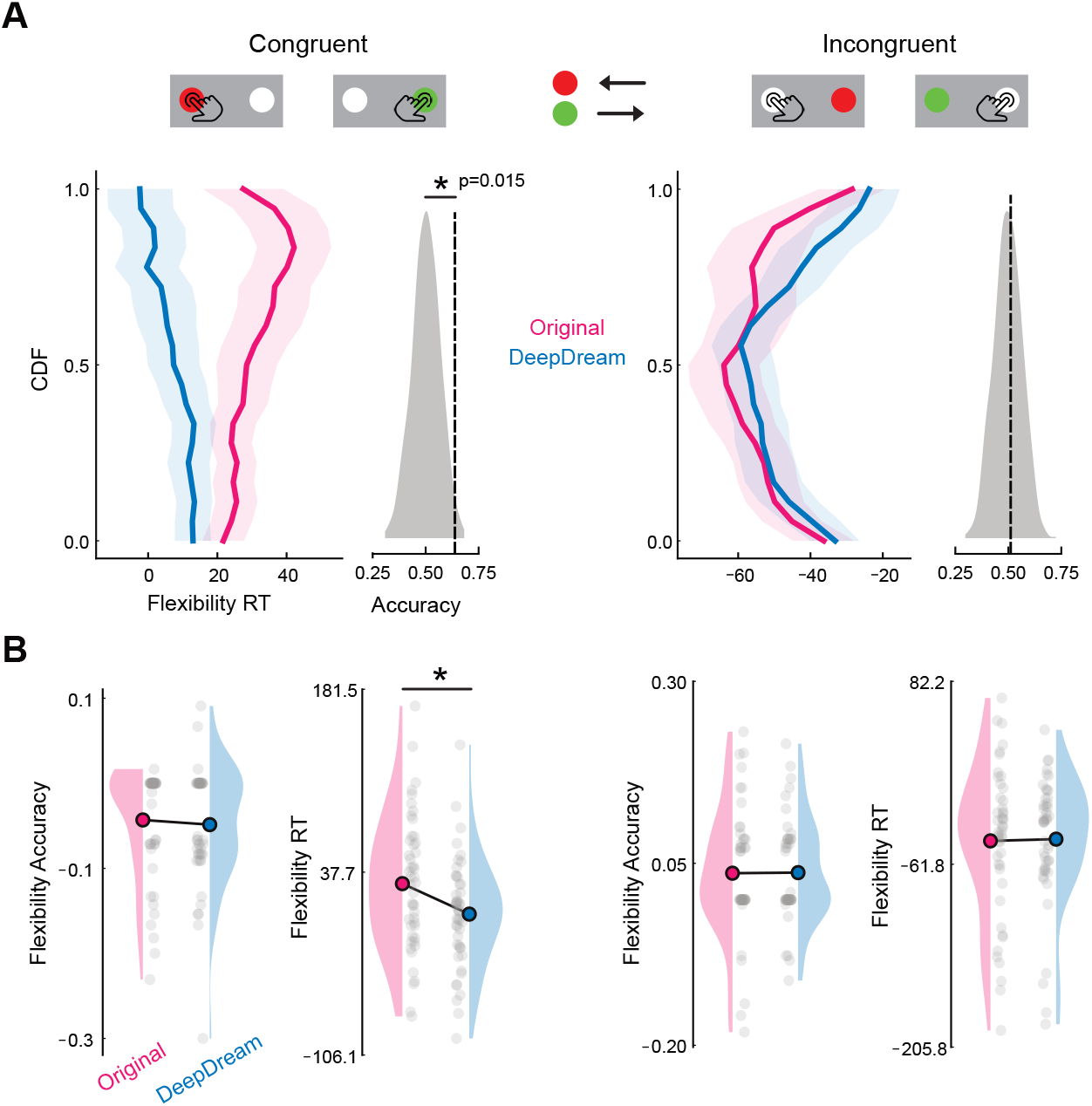
Immersive exposure to simulated hallucinations reduces the task switching costs. **A** Results from the Flexible Simon Task showing the flexibility score computed as the difference between the performance of the second block minus the first one. Blocks are distinguished by inverted stimulus-response mapping, leading to task switching costs when participants had to alternate the rule. Results are presented separately for congruent (left) and incongruent trials (right), showing the cumulative distribution function (CDF) of the reaction times (RT) for original (pink) and DeepDream (blue) on the y-axis and the flexibility score on the x-axis. On the side, there are the classification accuracy scores between conditions showed as a dotted vertical line and the grey distribution denote the empirical null distribution with permuted labels. Asterisks indicate statistical significance. **B** Raincloud plots showing the accuracy and RT flexibility scores for both conditions in congruent (left) and incongruent (right) trials. Colored circles indicate the mean of the distribution and grey dots are single participants. Asterisks indicate statistical significance.

Next, we asked whether the immersive exposure to DeepDream visual hallucinatory patterns modulated participants’ ability to generate linguistic associations (Fig. 3). Thus, we probed participants with a second task which was the Word Association Task ^22,23^ (Fig. 1B center) in which they had to associate a response word given a target word without overthinking (Fig. 3A). We leveraged word2vec deep language models ^24^ to extract word embedding vector representations of target and response words to compute a semantic distance score to investigate whether this measure differed across conditions. We found no difference in the semantic distances between target and response words (Fig. 3A left, p=0.501, d=0.013), nor between target and first response (Fig. 3A right, p=0.730, d=0.012) and the semantic distance between consecutive response words (Fig. 3A right, first and second response p=0.460, d=0.024, second and third response p=0.918, d=0.003). We also administered a third task to investigate linguistic associations beyond single words, namely the Story Continuation Task (Fig. 3B) ^25^. Here, participants were instructed to generate a story starting from an initial sentence (Fig. 1B center). To quantitatively assess the behavioral performance of participants at this task, we capitalized on recent transformer-based deep language models ^26^ to extract sentence-level embeddings of the response sentences. By computing the semantic distance between initial sentence and corresponding response sentence embeddings, we found no difference between OR and DD exposure (Fig. 3B left, p=0.952, d=0.007). Classification analyses between response sentence embeddings did not show a significant separation of the two conditions (Fig. 3B right, accuracy=51.2%, p=0.377). These results suggested that experiencing simulated psychedelic hallucinations did not modulate linguistic processing for word and sentence level associations.

**Figure 3:**
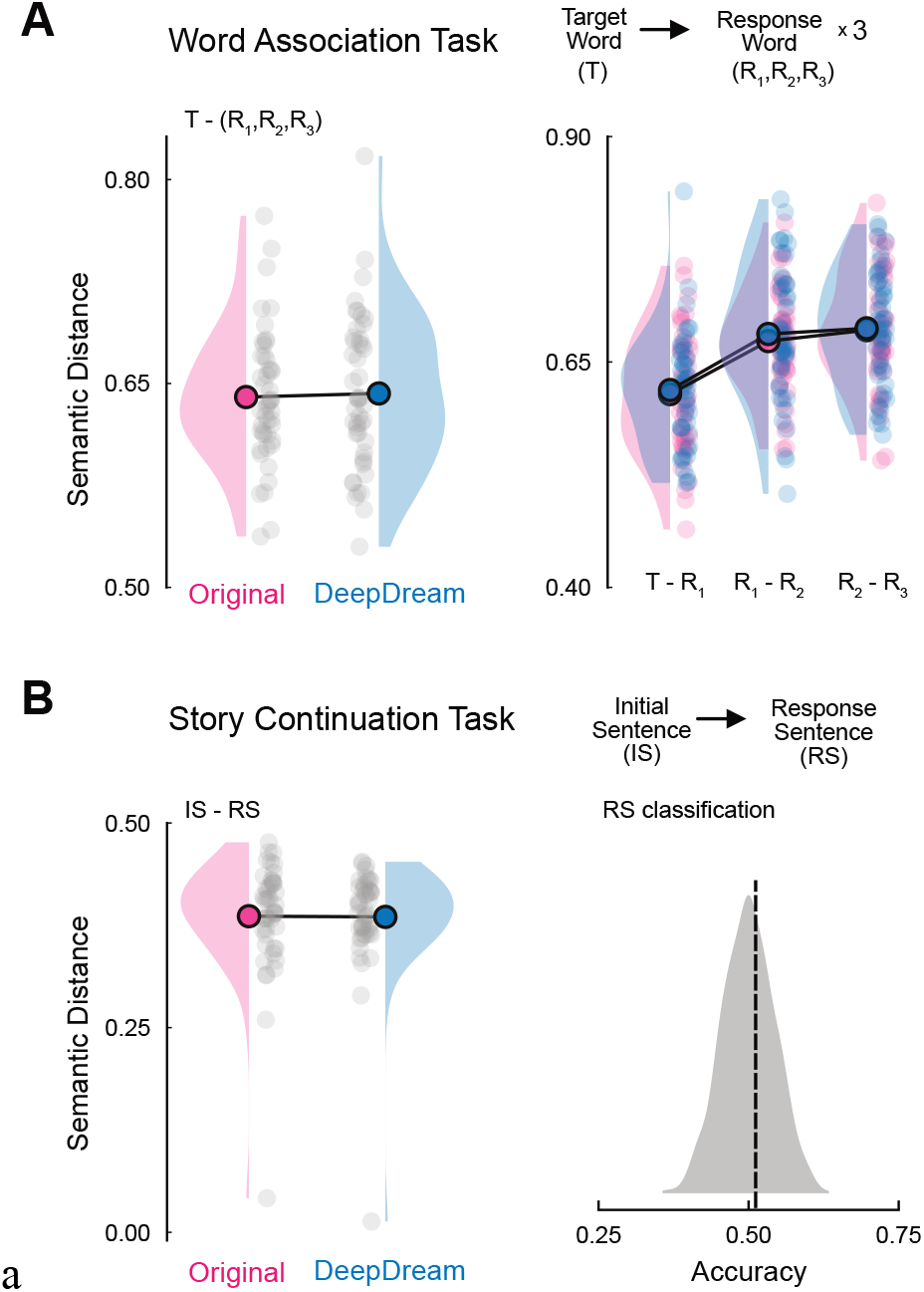
No evidence for simulated altered perception modulating linguistic stimuli-response associations. **A** Word Association Task with a target word and 3 response words associated. On the left, raincloud plots depicting the average semantic distance between target and all responses for original (pink) and DeepDream (blue). Colored circles indicate the mean of the distribution and grey dots are single participants. On the right, semantic distance between the target words and the first response and between consecutive responses. **B** Story continuation Task with an initial sentence (IS) to be continued with a response sentence (RS) to complete a story. On the left, raincloud plots showing the average semantic distance between the IS and the RS. Colored circles indicate the mean of the distribution and grey dots are single participants. On the right, classification accuracy scores by discriminating between the sentence embeddings of the two conditions RS indicated as a dotted vertical line and the grey distribution denoting the empirical null distribution with permuted labels.

Finally, we administered a fourth task, which we referred to as Creative Drawing Task (Fig. 1B right) ^27^, involving drawing a sketch starting from a visual cue, with the objective to be as creative as possible. Here, a creative drawing was operationalized as a sketch that was maximally divergent from the visual cue and at the same time was conveying a meaningful message through the whole sketch. Thus, we asked whether the exposure to DD lead to drawings having low- or high-level visual features different from OR. To answer this, we employed deep vision models ^28^ trained for image recognition to extract feature maps from the drawings and visual cues at different stages of the architectural hierarchy to disentangle low- and high-level features (Fig. 4A top). Thus, we measured relatedness of the visual cues and corresponding drawings in each condition by computing a similarity score between the feature maps extracted from the model (Fig. 4A bottom), which was the center kernel alignment (CKA) ^29^. We found that in early layers, encoding low-level features like spatial texture and edge distribution, OR and DD drawings did not differ in their CKA score (p=0.702, d=0.029). Importantly, we observed late layers, encoding object-level semantic information, showing a significantly difference between conditions (p=0.040, d=0.147). In particular, the similarity score of DD (average=0.208) was significantly less than OR (average=0.238), indicating that the exposure to simulated hallucinations lead participants to conceptually diverge from the visual cue significantly more than the naturalistic settings. To further test the hypothesis that DD exposure modulated high-level visual cognition, we leveraged current state-of-the-art multimodal deep vision-language models to extract rich text description from the drawings and investigate their semantic content (Fig. 4B top). We found this multimodal model being particularly effective at captioning non-trivial visual inputs such as drawings, as evidenced by a qualitative examination of the provided description given at some example drawings (Fig. 4B bottom left). Crucially, after converting these drawing related text captions into sentence embeddings, classification analysis revealed a significant discrimination of OR and DD based on the semantic content of the drawings (Fig. 4B bottom right, accuracy=58.0%, p=0.037). These results revealed how simulated psychedelic experience modulated high-level visual cognition, allowing participants to diverge from their visual inputs to generate semantically different visual abstract responses.

**Figure 4:**
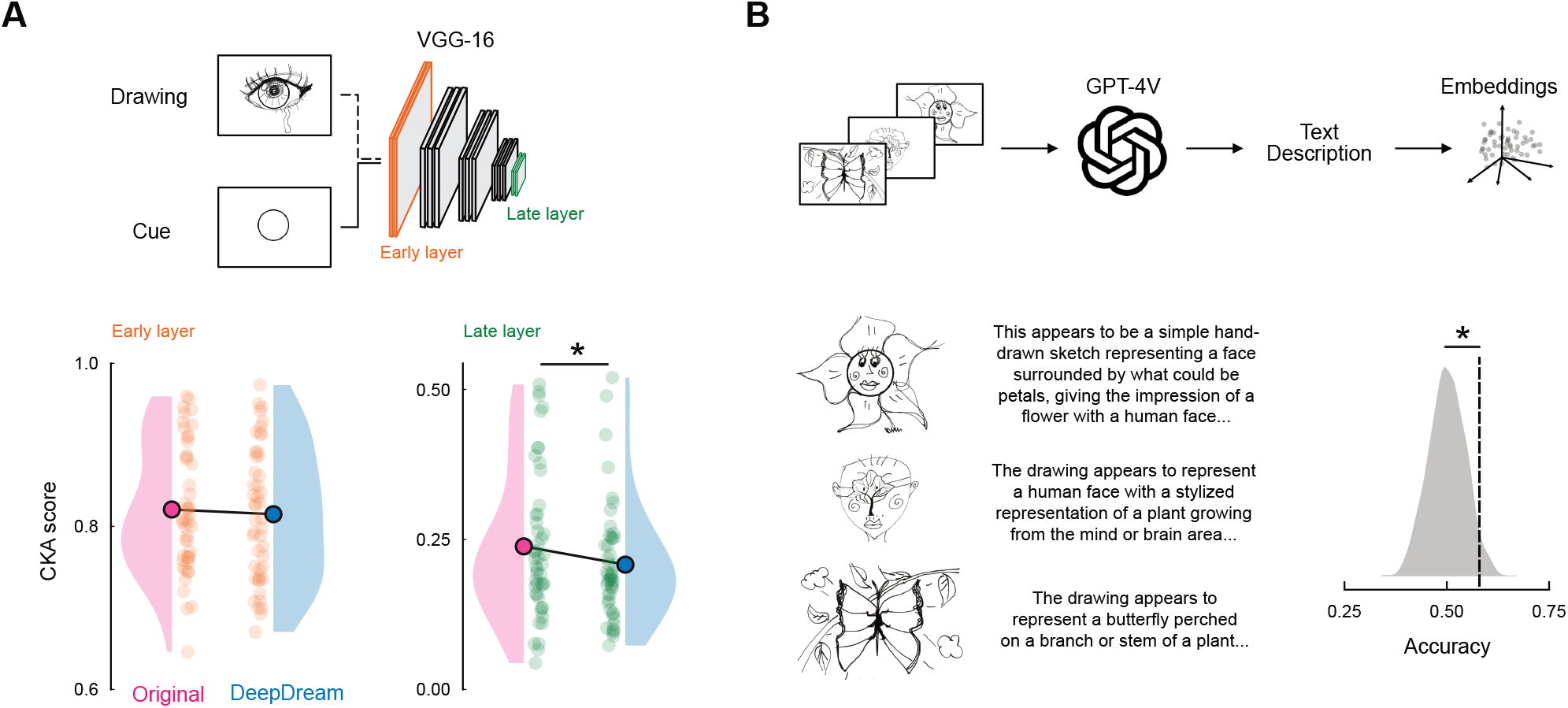
Immersive exposure to simulated psychedelic experience modulate visually grounded high-level cognition. **A** We passed the drawings collected from the Creative Drawing Task alongside their corresponding visual initial cue to the VGG-16 model. Then, we collected the feature maps from early (orange) and late (green) layers of the model, as shown in the top part of the panel. On the bottom, raincloud plots depicting the center kernel alignment (CKA) similarity scores between the cue and drawing responses for original (pink) and DeepDream (blue). Colored circles indicate the mean of the distribution and colored dots are individual participants. **B** We also probed GPT-4V by providing as input all the drawings and prompting it to deliver a rich textual description of the drawings (top). Then we converted this text description into sentence embeddings and used for classifying original and DeepDream conditions. On the bottom left, we showed some example drawings with their corresponding text description from the model, while on the bottom right we plotted the classification accuracy as a dotted vertical line with the grey distribution indicating the empirical null distribution with permuted labels. Asterisks indicate statistical significance.

## Discussion

Our investigation into the effects of perturbing low-level cognition through simulated visual hallucinations on high-level cognitive processes in humans has yielded significant insights into the relationship between perceptual and higher-order cognitive functions.

In the Flexible Simon Task ^20^, participants demonstrated reduced task-switching costs following the DeepDream video exposure compared to the naturalistic videos when instructed to invert the stimulus-response mapping. The effect was present only on congruent trials, probably due to a ceiling effect in the inherently more demanding incongruent trials. We interpreted this as evidence that executive functions such as cognitive flexibility was modulated by the altered perceptual manipulation. In particular, these findings not only replicate but also extend previous work by Rastelli et al. ^19^ demonstrating increased cognitive flexibility in a task measuring the ability to switch between stimulus-response mappings, as opposed to the inhibitory performance measured in their Stroop task.

Interestingly, our findings did not reveal significant effects of artificially altered phenomenology on linguistic cognitive abilities as measured by the Word Association Task (WAT) ^22,23^ and Story Continuation Task (SCT) ^25^, which measure the ability to make semantic associations at the word and sentence level, respectively. This apparent discrepancy with previous findings ^19^. may be attributed to our task instructions, which did not explicitly prompt creative responses or extensive exploration of the semantic space. In the WAT, participants were asked to associate a response word given a target word without overthinking, while similarly in the SCT, participants generated continuations of a given sentence to generate a simple story. Therefore, our instruction to participants to limit their exploration of the semantic space ^23^ may have been insufficient to reveal significant differences between the experimental conditions. Future research should employ tasks that incentivize broader semantic exploration, such as divergent thinking tasks, to further elucidate the impact of simulated visual hallucinations on language processing.

Our findings from the Creative Drawing Task ^27^ provided compelling evidence that artificially altered perception modulates high-level visual cognitive processes. Participants exposed to DeepDream videos produced semantically different and more abstract drawings compared to those who viewed original videos. This effect was quantified through higher dissimilarity between cue and response drawings in late layers of deep vision models and significant discriminability in sentence embeddings of drawing captions. This effect may be attributed to two factors: firstly, the explicit instruction to be “creative”, which encouraged participants to maximize divergence from the cue while maintaining meaningful content ^23,30^, and secondly, the visual nature of the task, which directly engaged the artificially altered sensory modality. Moreover, these findings align with reported effects of psychedelic drugs on visual imagery through modulation of functional connectivity between high-level and visual brain areas^31–33^.

Together, our results can be interpreted through the frameworks of predictive coding ^34–37^, the entropic brain hypothesis ^3,18^, and the REBUS (Relaxed Beliefs Under pSychedelics) theory ^4^.

Exposure to simulated hallucinations likely induced a state of higher entropy in the neural dynamics ^18^ and relaxed priors, similar to (but presumably milder than) psychedelic states. From a predictive coding perspective, the DeepDream-altered stimuli may have disrupted canonical predictive processes due to their out-of-distribution high-level visual features, leading to higher prediction errors and reconfiguration of sensory data generative models ^38,37^. The REBUS theory, which combines these frameworks, suggests that this exposure might have mimicked the relaxation of high-level priors that typically constrain cognition, allowing for more flexible updating of stimulus-response mappings and/or exploration of high-level visual concepts. These results support the notion that even brief exposure to stimuli that challenge generative predictive models of the sensory data can induce measurable changes in high-level cognitive processes such as cognitive flexibility and visual imagery, aligning with broader implications of these theories regarding the potential for perceptual alterations to induce more flexible states of cognition.

Crucially, our findings address a significant gap in the literature, which has extensively documented how high-level cognitive processes affect low-level perception ^39,40^, but offers limited evidence on the reverse influence. Our results underscore the strong interlinkage between bottom-up and top-down processes ^41,42^, demonstrating how perceptual alterations can induce measurable changes in high-level cognitive processes such as cognitive flexibility and visual imagery.

In conclusion, this study demonstrates that perturbing the visual perceptual phenomenology with simulated hallucinations significantly modulates high-level human cognition. These findings provide valuable insights into the interrelationship between bottom-up and top-down cognitive processes and open avenues for further scientific investigations into altered states of consciousness without pharmacological intervention. Future research should explore the duration and specificity of these effects, as well as their potential therapeutic applications in cognitive enhancement and mental health treatment.

## Acknowledgments

None

## Author Contributions

AG: Conceptualization, Methodology, Investigation, Data curation, Software, Formal analysis, Visualization, Writing – original draft, Writing – Review & Editing

CR: Conceptualization, Methodology, Investigation, Data curation, Software, Formal analysis, Visualization, Writing – original draft, Writing – Review & Editing

AU: Conceptualization, Methodology, Investigation, Writing – original draft, Writing – Review & Editing

GR: Conceptualization, Methodology, Supervision, Project administration, Funding acquisition, Writing – Review & Editing

## Data & Code Availability Statement

Data and custom code generated for this study are available from the corresponding authors upon reasonable request.

## Methods

### Participants

Forty-seven volunteers participated in the study (25 female and 22 male, average age of 28.42 years and a standard deviation of 10.17). All of them were native Italian speakers. No one of the participants reported having problems with their sight (normal or corrected-to-normal vision) and no one reported having neurological disorders or taking any neurological medications. Prior to the experiment, all participants provided written informed consent. All methods were approved by the University of Cattolica Milan, Human Research Ethics Committee (Protocol 69/24 March 2024). The whole procedure was realized in accordance with the Helsinki Declaration.

### Stimuli

The stimuli employed in this study were natural panoramic high-definition videos with 360° coverage recorded with a 360° panoramic camera (Insta360 X3) in Japan, including the audio track. The resolution of the videos was 5760 × 2880 (width and height) and were recorded at 30 frame per seconds (FPS), with an average bitrate of 200 Mbps. From these recordings, we selected 5 clips that represent everyday life in Tokyo (Shibuya Crossing and Akihabara), the deer park of Nara and a bus ride in Kyoto. The total duration of the sequence of these 5 clips was 10 minutes, with approximately 2 minutes for each clip. The order of presentation of the clips was fixed and we presented them to the participants one after the other without interruptions.

Then, we employed the DeepDream algorithm ^14,15,18,19^ to generate a perceptually altered version of the original video sequence. DeepDream is a computational technique that was initially conceived as a feature visualization procedure to investigate the hidden representations of pre-trained deep convolutional neural networks (CNN). It starts by forward passing an input image through the CNN up to a selected layer, with the objective function of maximizing its activation units. This is achieved via gradient ascent, which means adding the partial derivatives (gradients) of the objective function with respect to the input image to the image itself. This optimization procedure alters the image by accentuating the visual features that are preferred to that hidden layer, a process that has been referred to as “algorithmic pareidolia” ^19^. What stands out about this generative procedure is that the images produced frequently exhibit a distinct “hallucinatory” character, showing an intuitive resemblance to the diverse array of psychedelic visual experiences documented in literature ^14,19,43^. Following previous studies applying DeepDream to videos for artificial psychedelic research ^14,19^, we used dense optical flow estimation and alpha blending to perceptually stabilize the generation process of each frame. The patterns resembling hallucinations in the previous frame, generated in regions where optical flow is observed, are combined with the current frame using a specific blending ratio before applying DeepDream. The blending ratio was separated for parts of the frame with low and high estimated flow. This adjustment ensures that areas with lesser optical flow are not overwhelmed by the more intense blending ratios designated for areas with significant optical flow. In this study, we set a threshold of 6 to divide the frame pixels into low and high flow and pplied a bending ration of 0.9 to the high flow and 0.1 to the low flow. Optical flow was estimated using the Farnebäck algorithm ^44^ with a pyramid scale set as 0.5 with 5 levels, an averaging window size of 13 with 10 iterations per level, 5 pixel neighbours and a Gaussian kernel with a standard deviation of 1.1. We also used two pretrained GoogleNet (InceptionV3) CNN ^45^, one trained on Imagenet1K ^46^ and one on Places365 ^47^. The former one we used to “hallucinate” 3 clips and the latter for the remaining 2 clips. For both models, we selected the “inception_4d_pool” layer and used 4 octaves with an octave scale parameter of 1.2, jitter as 32 pixels and video frames resolution was rescaled by a factor of 2. Optimization was carried out by gradient ascent with normalized gradients with 15 iterations and a learning rate of 1.5.

### Procedure

Participants were welcomed in the dedicated experimental setting, gave consent and we explained all the information about the experiment. Then the head-mounted display (HMD, here a Meta Quest 2) was adjusted for their head size. The experiment consisted of two conditions, original (OR) and DeepDream (DD), in which participants were exposed to panoramic videos in VR, with a total duration of 10 min. The order of conditions was counterbalanced across participants. After the HMD was fitted, participants, while sitting, were encouraged to explore the video sequence. After the exposure to each condition, participants performed four behavioural tasks, namely the Flexible Simon task (FST), the Word Association task (WAT), the Story Continuation task (SCT) and the Creative Drawing task (CDT). All tasks were performed always in the same presented order, the first three via a computer screen implemented in Psytoolkit ^48^ and the CDT on an iPad with a tablet pen.

The FST consisted of a modified version of the Simon Test ^20^. This task can study the interference between spatially located stimuli and motor responses to assess cognitive flexibility. The stimuli consisted of a red or green circle located in the right part or the left part of the computer screen. If, for example, the left circle was red the right one was left white. Participants were instructed to press on the keyboard the key A (positioned on the left side of the keyboard) with their left index finger whenever they saw the red circle, independently of the position of the circle. Similarly, they were instructed to press the key L (on the right of the keyboard) with their right index finger whenever they saw the green circle. They were also instructed to press as fast as possible. A total of 32 trials were presented in randomized order, whose 50% matched the position of the circle (left or right) on the screen with the positions of the keys A or L (congruent stimuli), while in the remaining 50%, the position of the stimulus and the position of the target key were unmatching (incongruent trials). After this block, participants were instructed to invert the mapping of stimuli-response and we administered other 32 trials with the same structure as the first block.

The WAT probed participants’ semantic memory by asking them to generate associations between words representing concrete objects ^22,23^. We instructed participants to generate 3 response words given a target word according to a set of instructions involving the constrain of responses being only concrete nouns, no names of people or famous objects and no technical words or jargon. We also asked them to respond as fast as possible from the onset of the presentation of the target word on the screen. The target words were 10 and were extracted from 5 different semantic categories. These target words were divided into two sets and were counterbalanced across participants for their assignation to each experimental condition, consisting of the following words (in Italian): “Dog”, “Rabbit”, “Water”, “Stone”, “Chair”, “Bathtub”, “Hand”, “Mouth”, “School” and “Shop” for the first target word list and “Sand”, “Fire”, “Hospital”, “Gym”, “Nose”, “Head”, “Cat”, “Horse”, “Table” and “Pencil” for the second target word list. The order of presentation of the target words was randomized.

The SCT consisted of requiring participants to complete an unfinished sentence, with the goal to generate a simple story ^25^. Participants were instructed to complete the story within 60 seconds. The initial sentence was presented for the whole time they were generating the response. There were 2 starting sentences presented for each condition and we divided them in two sets and counterbalanced across participants as above. These starting sentences were the following (in Italian): “At a certain point, he found himself in the middle of a dense forest, disoriented and with no memory of how he had ended up there…” and “There once was a shrimp that…” for the first set, while “Outside, it was raining, and a cat was…” and “They had always been very unlucky in what they did when one day…” were assigned to the second set.

The CDT consisted of asking participants of drawing a sketch starting from a blank page with a visual cue in the center ^27^. Participants were instructed to generate a drawing as creative as possible, which was operationalized as attempting to conceptually maximize the divergence from the cue and at the same time trying to convey a meaningful message through the whole sketch. They were required to generate a drawing within 120 seconds and 2 visual cues were presented per condition. As above, we created two sets of visual cues and counterbalanced across participants for the assignment to each condition and the visual cues were a “circle” and a “man” for the first set and a “square” and a “tree” for the second set.

### Data analysis

We started the data analysis of the behavioral data with the FST data. The data consisted of the accuracy and reaction times (RT) of participants for each trial. We preprocessed these data by removing outlier responses given the RT distribution for each participant. Specifically, we discarded the trials with a median absolute deviation (MAD) being ±2.5 times the median value. Then, we defined a flexibility score by subtracting the average performance of the first block (with inverted stimulus-response mapping) from the average performance of the second block. This allowed us to investigate the switching cost of the participants given the change of the mapping rule ^21^. Thus, the greater this flexibility score is, the greater are the costs of switching for the participant. We applied this score to the accuracy, the empirical cumulative distribution function (CDF) of the RT and the mean RT separately for the congruent and incongruent trials and the experimental conditions. We also compared CDF between conditions using multivariate classification analyses (see statistical analysis section below).

Then, we analyzed the WAT data using a distributional semantics approach, grounded on the assumption that semantically similar words appear in similar contexts ^49^. We extracted word embeddings (i.e., vectorial representations of words) for all target and response words using a pretrained word2vec model ^24^. This model is a deep neural network (DNN) that employ a sliding window approach to move through text corpora, namely continuous bag of words (CBOW). CBOW is trained to maximize the probability of the target word by looking at the context, where the context is represented as a bag of words contained in a fixed size window around the target word. A word embedding vector representation is then extracted from the hidden layer of CBOW. Word2vec models have already been shown to correlate highly with human judgments of semantic relatedness ^50^. The model was trained using fastText on a concatenation of the Common Crawl and Wikipedia ^51^. Thus, we computed the semantic distance between all pairs of target and response words and between consecutive response words as the cosine distance between the related word embeddings ^23,52^.

Next, we analyzed the SCT data by capitalizing on current state-of-the-art natural language processing (NLP) transformer models, namely the Bidirectional Encoder Representations from Transformers (BERT) model ^26^. Trained on extensive text corpora, BERT is able to capture complex syntactic and semantic dependencies between words, taking into account the full context of a sentence by analyzing words bidirectionally (i.e., in both forward and backward directions). The underlying architecture is based on the Transformer ^53^, a DNN architecture that operates on the mechanisms of attention, enabling the model to focus on various elements of the text. We used an instance of the model from HugginFace ^54^, namely the Italian BERTino ^55^, a distilBERT model ^56^ trained on the Paisa and ItWaC corpora ^57^. Thus, we forward passed through the model the tokenized version of the initial sentences and initial plus response sentences. We collected the last hidden representation layer of the classification token (CLS), which encodes a context-dependent sentence-level representation of the text input with a dimensionality of 768. Then, we computed the semantic distance as above between the initial and initial plus response sentence embeddings and compared the sentence embeddings between conditions using classification analysis.

Lastly, we analyzed the CDT data by means of deep visual and multimodal models. We employed the VGG-16 model ^28^ to quantitatively assess that the drawings differed between original and DeepDream exposure in their level of visual abstraction. VGG-16 is a CNN trained on Imagenet1K ^46^ and it is widely used not only because it achieved good performance in image recognition tasks but also since there is abundant evidence that it has learned visual representations that resemble visual object representations in the human brain ^58–61^, specifically in the ventral stream ^62^. We adopted the VGG-16 implementation from the torchvision library ^63^ and forward passed the drawings and their corresponding visual cues to the model, extracting early and late feature maps layers. We used this methodology since it is known that these CNN models emulate the sequential extraction of visual information from low-level features in the initial layers to high-level semantic content in more advanced layer in the hierarchy ^64–67^. As the early layer representation, we selected the first convolutional layer after the rectified linear unit (ReLU) activation function (“features.1” using the torchvision nomenclature), while as the late layer we used the last convolutional layer after the 2D max pooling operation (“features.30” using the torchvision nomenclature). Center Kernel Alignment (CKA) ^29^ with a linear kernel was used as a similarity score of the models’ layer representations between the drawings and their corresponding visual cues. In addition to this pipeline employing deep visual models, we also leveraged current state-of-the-art deep multimodal vision-language models to further test the semantic content of the drawing between conditions. We used the generative pretrained transformer 4 vision (GPT-4V) model from OpenAI application programming interface (API) ^68^, arguably one of the most highly performing multimodal models in visual understanding and image captioning tasks currently available ^69–71^. For each drawing, we asked GPT-4V to provide an image caption (i.e., a text description) using the following prompt: “This is a drawing. Please, describe what the author wants to represent.”. We also set the output to have a maximum of 300 tokens. Once obtained the captions, we used another sentence embedding model from the same API, namely the “text-embedding-3-small”, to convert the textual description into a sentence embedding vector with a dimensionality of 1536. We discarded 23 drawing descriptions (9 original) since the GPT-4V model did not provide an appropriate output. We used these embeddings to perform a classification analysis, by compensating the class imbalance with copying random instances from the minority class to match the numerosity of the majority class.

### Statistical analysis

Statistical analysis was carried out using linear mixed-effect models (LMM) whenever it was required a univariate comparison between the original and DeepDream condition. We used LMM to account for the nested data structure given by the fact that we used all the trials from all participants to estimate the effect at the group level. The model was generally specified with the dependent variable being the measure calculated for each analysis, the fixed effect was the condition factor and the participant identity was modeled as a random effects to account for the variation across individuals. In addition, whenever it was required a multivariate comparison such as in the CDF of the reaction times or the embedding vectors, we performed a binary classification analysis between the original and DeepDream conditions using a support vector machine (SVM) with a radial basis function (RBF) as kernel ^72^. We used a 3-fold cross validation to estimate the out-of-distribution performance of the model and avoid overfitting. Z-scoring was applied by computing the sample mean and standard deviation on the train set and then applying them to the test set to avoid possible confounders. When classifying the conditions using embedding vectors, we additionally performed principal component analysis (PCA) to reduce the dimensionality of the data to 20 principal components, again estimating the loadings on the training set and using them to project in the latent space the test data to avoid data leakage. Accuracy was used as metric performance. To statistically validate the model performance, we performed a permutation test by randomly shuffling the labels assigned to each sample and rerun the classification analysis with 1000 permutations. P-values were computed as the proportion of permuted classification accuracies that exceeded the actual model accuracy.

## References

1. Carhart-Harris, R. L. et al. Neural correlates of the psychedelic state as determined by fMRI studies with psilocybin. Proceedings of the National Academy of Sciences 109, 2138–2143 (2012).

2. Carhart-Harris, R. L. et al. Neural correlates of the LSD experience revealed by multimodal neuroimaging. Proceedings of the National Academy of Sciences of the United States of America 113, 4853–4858 (2016).

3. Carhart-Harris, R. L. et al. The entropic brain: a theory of conscious states informed by neuroimaging research with psychedelic drugs. Frontiers in Human Neuroscience 8, 20 (2014).

4. Carhart-Harris, R. L. & Friston, K. J. REBUS and the Anarchic Brain: Toward a Unified Model of the Brain Action of Psychedelics. Pharmacol Rev 71, 316–344 (2019).

5. Gründer, G. et al. Treatment with psychedelics is psychotherapy: beyond reductionism. The Lancet Psychiatry 11, 231–236 (2024).

6. Carhart-Harris, R. et al. Trial of Psilocybin versus Escitalopram for Depression. N Engl J Med 384, 1402–1411 (2021).

7. Scott, G. & Carhart-Harris, R. L. Psychedelics as a treatment for disorders of consciousness. Neuroscience of Consciousness 2019, niz003 (2019).

8. Carhart-Harris, R. L. et al. Psilocybin with psychological support for treatment-resistant depression: an open-label feasibility study. The Lancet Psychiatry 3, 619–627 (2016).

9. Oxman, T. E., Rosenberg, S. D., Schnurr, P. P., Tucker, G. J. & Gala, G. The language of altered states. J Nerv Ment Dis 176, 401–408 (1988).

10. Tagliazucchi, E., Carhart-Harris, R., Leech, R., Nutt, D. & Chialvo, D. R. Enhanced repertoire of brain dynamical states during the psychedelic experience. Human Brain Mapping 35, 5442–5456 (2014).

11. Bayne, T. & Carter, O. Dimensions of consciousness and the psychedelic state. Neurosci Conscious 2018, niy008 (2018).

12. Shinozuka, K. et al. LSD flattens the hierarchy of directed information flow in fast whole-brain dynamics. bioRxiv (2024) doi:10.1101/2024.04.25.591130.

13. Preller, K. H. et al. Effective connectivity changes in LSD-induced altered states of consciousness in humans. Proceedings of the National Academy of Sciences 116, 2743–2748 (2019).

14. Suzuki, K., Roseboom, W., Schwartzman, D. J. & Seth, A. K. A deep-dream virtual reality platform for studying altered perceptual phenomenology. Scientific Reports 7, 1–11 (2017).

15. Mordvintsev, A., Olah, C. & Tyka, M. Inceptionism: Going Deeper into Neural Networks. Google Research Blog Available at: http://googleresearch.blogspot.co.uk/2015/06/inceptionism-going-deeper-into-neural.html. http://googleresearch.blogspot.co.uk/2015/06/inceptionism-going-deeper-into-neural.html (2015).

16. Krizhevsky, A., Sutskever, I. & Hinton, G. E. ImageNet Classification with Deep Convolutional Neural Networks. in Advances in Neural Information Processing Systems vol. 25 (Curran Associates, Inc., 2012).

17. LeCun, Y., Bengio, Y. & Hinton, G. Deep learning. Nature 521, 436–444 (2015).

18. Greco, A., Gallitto, G., D’Alessandro, M. & Rastelli, C. Increased Entropic Brain Dynamics during DeepDream-Induced Altered Perceptual Phenomenology. Entropy 23, 839 (2021).

19. Rastelli, C., Greco, A., Kenett, Y. N., Finocchiaro, C. & De Pisapia, N. Simulated visual hallucinations in virtual reality enhance cognitive flexibility. Scientific reports 12, 4027 (2022).

20. Simon, J. R. & Rudell, A. P. Auditory SR compatibility: the effect of an irrelevant cue on information processing. Journal of applied psychology 51, 300 (1967).

21. Uddin, L. Q. Cognitive and behavioural flexibility: neural mechanisms and clinical considerations. Nature Reviews Neuroscience 22, 167–179 (2021).

22. Vivas, L., Manoiloff, L., García, A. M., Lizarralde, F. & Vivas, J. Core Semantic Links or Lexical Associations: Assessing the Nature of Responses in Word Association Tasks. J Psycholinguist Res 48, 243–256 (2019).

23. Rastelli, C., Greco, A., De Pisapia, N. & Finocchiaro, C. Balancing novelty and appropriateness leads to creative associations in children. PNAS nexus 1, pgac273 (2022).

24. Mikolov, T., Sutskever, I., Chen, K., Corrado, G. S. & Dean, J. Distributed representations of words and phrases and their compositionality. Advances in neural information processing systems 26, (2013).

25. Holmes, V. M. Sentence Planning in a Story Continuation Task. Lang Speech 27, 115–134 (1984).

26. Devlin, J., Chang, M.-W., Lee, K. & Toutanova, K. BERT: Pre-training of Deep Bidirectional Transformers for Language Understanding. Preprint at 10.48550/arXiv.1810.04805 (2019).

27. Fan, J. E., Bainbridge, W. A., Chamberlain, R. & Wammes, J. D. Drawing as a versatile cognitive tool. Nat Rev Psychol 2, 556–568 (2023).

28. Simonyan, K. & Zisserman, A. Very Deep Convolutional Networks for Large-Scale Image Recognition. Preprint at http://arxiv.org/abs/1409.1556 (2015).

29. Kornblith, S., Norouzi, M., Lee, H. & Hinton, G. Similarity of Neural Network Representations Revisited. Preprint at 10.48550/arXiv.1905.00414 (2019).

30. Rastelli, C. et al. Neural dynamics of semantic control underlying generative storytelling. bioRxiv (2024).

31. Kometer, M., Cahn, B. R., Andel, D., Carter, O. L. & Vollenweider, F. X. The 5-HT2A/1A agonist psilocybin disrupts modal object completion associated with visual hallucinations. Biol Psychiatry 69, 399–406 (2011).

32. de Araujo, D. B. et al. Seeing with the eyes shut: neural basis of enhanced imagery following ayahuasca ingestion. Human Brain Mapping 33, 2550–2560 (2012).

33. Stoliker, D. et al. Neural mechanisms of psychedelic visual imagery. Mol Psychiatry 1–8 (2024) doi:10.1038/s41380-024-02632-3.

34. Rao, R. P. & Ballard, D. H. Predictive coding in the visual cortex: a functional interpretation of some extra-classical receptive-field effects. Nature neuroscience 2, 79–87 (1999).

35. Friston, K. The free-energy principle: a unified brain theory? Nature Reviews Neuroscience 11, 127–138 (2010).

36. Clark, A. Whatever next? Predictive brains, situated agents, and the future of cognitive science. Behavioral and brain sciences 36, 181–204 (2013).

37. Greco, A., Moser, J., Preissl, H. & Siegel, M. Predictive learning shapes the representational geometry of the human brain. bioRxiv 2024–03 (2024).

38. Bonna, K., Hulme, O. J., Meder, D., Duch, W. & Finc, K. Brain network reconfiguration during prediction error processing. bioRxiv 2023.07.14.549018 (2023) doi:10.1101/2023.07.14.549018.

39. Lupyan, G. & Ward, E. J. Language can boost otherwise unseen objects into visual awareness. Proceedings of the National Academy of Sciences 110, 14196–14201 (2013).

40. Francken, J. C., Kok, P., Hagoort, P. & de Lange, F. P. The behavioral and neural effects of language on motion perception. Journal of Cognitive Neuroscience 27, 175–184 (2015).

41. Goldstone, R. L. & Barsalou, L. W. Reuniting perception and conception. Cognition 65, 231–262 (1998).

42. Barsalou, L. W. Grounded Cognition. Annual Review of Psychology 59, 617–645 (2008).

43. McKenna, D. J. Clinical investigations of the therapeutic potential of ayahuasca: rationale and regulatory challenges. Pharmacology & therapeutics 102, 111–129 (2004).

44. Farnebäck, G. Two-Frame Motion Estimation Based on Polynomial Expansion. in Image Analysis (eds. Bigun, J. & Gustavsson, T.) vol. 2749 363–370 (Springer Berlin Heidelberg, Berlin, Heidelberg, 2003).

45. Szegedy, C. et al. Going deeper with convolutions. in Proceedings of the IEEE conference on computer vision and pattern recognition 1–9 (2015).

46. Deng, J. et al. Imagenet: A large-scale hierarchical image database. in 2009 IEEE conference on computer vision and pattern recognition 248–255 (Ieee, 2009).

47. Zhou, B., Lapedriza, A., Khosla, A., Oliva, A. & Torralba, A. Places: A 10 million image database for scene recognition. IEEE transactions on pattern analysis and machine intelligence 40, 1452–1464 (2017).

48. Stoet, G. PsyToolkit: A Novel Web-Based Method for Running Online Questionnaires and Reaction-Time Experiments. Teaching of Psychology 44, 24–31 (2017).

49. Boleda, G. Distributional Semantics and Linguistic Theory. Annu. Rev. Linguist. 6, 213–234 (2020).

50. Mandera, P., Keuleers, E. & Brysbaert, M. Explaining human performance in psycholinguistic tasks with models of semantic similarity based on prediction and counting: A review and empirical validation. Journal of Memory and Language 92, 57–78 (2017).

51. Grave, E., Bojanowski, P., Gupta, P., Joulin, A. & Mikolov, T. Learning Word Vectors for 157 Languages. Preprint at http://arxiv.org/abs/1802.06893 (2018).

52. Heinen, D. J. P. & Johnson, D. R. Semantic distance: An automated measure of creativity that is novel and appropriate. Psychology of Aesthetics, Creativity, and the Arts 12, 144–156 (2018).

53. Vaswani, A. et al. Attention is all you need. Advances in neural information processing systems 30, (2017).

54. Wolf, T. et al. HuggingFace’s Transformers: State-of-the-art Natural Language Processing. Preprint at 10.48550/arXiv.1910.03771 (2020).

55. Muffo, M. & Bertino, E. BERTino: an Italian DistilBERT model. Preprint at 10.48550/arXiv.2303.18121 (2023).

56. Sanh, V., Debut, L., Chaumond, J. & Wolf, T. DistilBERT, a distilled version of BERT: smaller, faster, cheaper and lighter. Preprint at 10.48550/arXiv.1910.01108 (2020).

57. Lyding, V. et al. The PAISÀ Corpus of Italian Web Texts. in Proceedings of the 9th Web as Corpus Workshop (WaC-9) (eds. Bildhauer, F. & Schäfer, R.) 36–43 (Association for Computational Linguistics, Gothenburg, Sweden, 2014). doi:10.3115/v1/W14-0406.

58. Storrs, K. R., Kietzmann, T. C., Walther, A., Mehrer, J. & Kriegeskorte, N. Diverse Deep Neural Networks All Predict Human Inferior Temporal Cortex Well, After Training and Fitting. Journal of Cognitive Neuroscience 33, 2044–2064 (2021).

59. Schrimpf, M. et al. Brain-score: Which artificial neural network for object recognition is most brain-like? BioRxiv 407007 (2018).

60. Schrimpf, M. et al. Integrative Benchmarking to Advance Neurally Mechanistic Models of Human Intelligence. Neuron 108, 413–423 (2020).

61. Singer, J. J. D., Cichy, R. M. & Hebart, M. N. The Spatiotemporal Neural Dynamics of Object Recognition for Natural Images and Line Drawings. J. Neurosci. 43, 484–500 (2023).

62. Güçlü, U. & Van Gerven, M. A. Deep neural networks reveal a gradient in the complexity of neural representations across the ventral stream. Journal of Neuroscience 35, 10005–10014 (2015).

63. maintainers, T. & contributors. TorchVision: PyTorch’s Computer Vision library. GitHub repository GitHub (2016).

64. Cadieu, C. F. et al. Deep neural networks rival the representation of primate IT cortex for core visual object recognition. PLoS computational biology 10, e1003963 (2014).

65. Yamins, D. L. K. et al. Performance-optimized hierarchical models predict neural responses in higher visual cortex. Proc. Natl. Acad. Sci. U.S.A. 111, 8619–8624 (2014).

66. Yamins, D. L. & DiCarlo, J. J. Using goal-driven deep learning models to understand sensory cortex. Nature neuroscience 19, 356–365 (2016).

67. Zeiler, M. D. & Fergus, R. Visualizing and Understanding Convolutional Networks. Preprint at 10.48550/arXiv.1311.2901 (2013).

68. GPT-4V(ision) system card. https://openai.com/research/gpt-4v-system-card.

69. Wu, Y. et al. An Early Evaluation of GPT-4V(ision). Preprint at 10.48550/arXiv.2310.16534 (2023).

70. Yang, Z. et al. The dawn of lmms: Preliminary explorations with gpt-4v (ision). arXiv preprint 2309.17421 9, 1 (2023).

71. Zhang, C. & Wang, S. Good at captioning, bad at counting: Benchmarking GPT-4V on Earth observation data. Preprint at 10.48550/arXiv.2401.17600 (2024).

72. Cortes, C. & Vapnik, V. Support-vector networks. Mach Learn 20, 273–297 (1995).

